# *Agrobacterium*-Mediated Transient Transformation of *Marchantia* Liverworts

**DOI:** 10.1101/2021.05.14.444154

**Authors:** Hidekazu Iwakawa, Katharina Melkonian, Titus Schlüter, Ryuichi Nishihama, Hiroyasu Motose, Hirofumi Nakagami

## Abstract

*Agrobacterium*-mediated transient gene expression is a rapid and useful approach for characterizing functions of gene products *in planta*. However, the practicability of the method in the model liverwort *Marchantia polymorpha* has not yet been thoroughly described. Here we report a simple and robust method for *Agrobacterium*-mediated transient transformation of *Marchantia* thalli and its applicability. When thalli of *M. polymorpha* were co-cultured with *Agrobacterium tumefaciens* carrying *GUS* genes, GUS staining was observed primarily in assimilatory filaments and rhizoids. GUS activity was detected 2 days after infection and saturated 3 days after infection. We were able to transiently co-express fluorescently tagged proteins with proper localizations. Furthermore, we demonstrate that our method can be used as a novel pathosystem to study liverwort-bacteria interactions. We also provide evidence that air chambers support bacterial colonization.

## Introduction

*Marchantia polymorpha*, or the common liverwort, is a member of the non-vascular plant lineage and has been established as a new experimental model in recent years (Bowman et al. 2016; Bowman et al. 2017; Ishizaki et al. 2016). The genome of *M. polymorpha* subspecies *ruderalis* strain Takaragaike-1 (Tak-1) has been sequenced (Bowman et al. 2017), and a variety of molecular genetic tools and techniques have been established for the species (Boehm et al. 2016; Flores-Sandoval et al. 2016; Ishizaki et al. 2016; Nishihama et al. 2016; Shimamura 2016b; Sugano et al. 2018; Tanaka et al. 2016). The Takaragaike accessions, Tak-1 and Tak-2, were isolated in Kyoto, Japan, and have been widely utilized for molecular studies (Okada et al. 2000). Besides Tak-1 and Tak-2, an Australian population of *M. polymorpha* isolated from a field location near Melbourne, Victoria (Flores-Sandoval et al. 2015), and *M. polymorpha* accession BoGa obtained from the Botanical Garden of Osnabrück, Germany (Althoff et al. 2014; Buschmann et al. 2016), have been utilized. Most thalloid liverworts, including *Marchantia* species, are known to form mycorrhizal associations (Ligrone et al. 2007; Russell and Bulman 2005). However, similar to mosses, *M. polymorpha* have most likely lost genes involved in symbiosis, and, therefore, cannot form beneficial associations with mycorrhizal fungi (Bowman et al. 2017; Ligrone et al. 2007). For this reason, another *Marchantia* species, *M. paleacea*, which retains symbiotic ability (Humphreys et al. 2010; Radhakrishnan et al. 2020; Rimington et al. 2018), is gaining in popularity as a model to study symbiosis in liverworts, although molecular genetic tools for this species are as yet limited.

Transient gene expression or silencing is a rapid and useful approach to characterize gene or protein functions *in planta*, but has yet been scarcely reported for the liverworts. To date, several approaches based on electroporation, polyethylene glycol (PEG), viruses, biolistics, or the *Agrobacterium*, have been established to deliver foreign genes into plant cells (Fromm et al. 1985; Kapila et al. 1997; Klein et al. 1987; Negrutiu et al. 1987; Takamatsu et al. 1987). Recently, Westermann *et al*. reported that a particle bombardment method can be used to transiently transform epidermal cells of *Marchantia* thalli (Westermann et al. 2020). *Agrobacterium tumefaciens* was originally isolated as a tumorigenic pathogen causing crown gall disease in a wide range of eudicots (Chilton et al. 1977). This ability is dependent on the introduction of transfer DNA (T-DNA) into the plant genome, and elucidation of the mechanism of T-DNA transfer has enabled scientists to engineer *A. tumefaciens* that transfer selected genes into plant cells without causing disease. *Agrobacterium*-mediated transformation has thus been widely applied in plant research. The *Agrobacterium*-mediated stable transformation of plant cells involves eight crucial steps (Gelvin 2010): 1) *Agrobacterium* attachment and biofilm formation at the plant cell surface, 2) injection of T-strand and virulent effector (Vir) proteins into the plant cell via a type IV secretion system, 3) formation of a T-complex, composed of the T-strand and Vir proteins inside the plant cell, 4) super T-complex formation by association with plant proteins, such as VirE2-binding protein (VIP1) and importin α, 5) translocation of the super T-complex into the nucleus, 6) chromatin targeting of the super T-complex and dissociation of the proteins, 7) integration of the T-strand into the plant genome, and 8) expression of transgenes encoded on the T-DNA. Since the sixth and seventh steps in which the T-DNA is integrated into the plant genome are not prerequisites for transgene expression, the *Agrobacterium* can also be utilized to transiently express selected genes in plant cells (Gelvin 2010).

Plant cells can recognize access of *Agrobacterium* through their plasma membrane localized microbe-associated molecular pattern (MAMP) receptors, which activate pattern-triggered immunity (PTI) and thereby restrict *Agrobacterium*-mediated transformation (Rosas-Díaz et al. 2017; Tsuda et al. 2012; Zhu et al. 2017; Zipfel et al. 2006). Suppression of PTI by use of the effector protein AvrPto from the bacterial pathogen *Pseudomonas syringae* could improve efficiency of *Agrobacterium*-mediated transient transformation in *Arabidopsis thaliana* (Tsuda et al. 2012). Additionally, salicylic acid (SA)-mediated immune responses are known to restrict *Agrobacterium*-mediated transient transformation. Efficient transient expression was observed in *A. thaliana sid2, npr1*, and NahG plants that have a defect in SA pathways (Rosas-Díaz et al. 2017; Tsuda et al. 2012; Zhu et al. 2017). In contrast, *A. thaliana coi1* mutant, lacking a jasmonate (JA) receptor, further restricted the transient expression, which can be explained by SA-JA antagonism (Rosas-Díaz et al. 2017).

The hemi-biotrophic pathogenic bacterium *Pseudomonas syringae* pv. *tomato* DC3000 (*Pto* DC3000), causal agent of tomato bacterial speck disease, has been widely used as a model pathogen to understand the plant immune system (Xin and He 2013). Recently, it was shown that *Pto* DC3000 can infect and cause disease in *M. polymorpha* in an effector-dependent manner (Gimenez-Ibanez et al. 2019). The effector proteins AvrPto or AvrPtoB were transiently expressed in *M. polymorpha* thalli by using *A. tumefaciens*, and the expression of effectors was shown to suppress the marker gene expression induced by crude extracts from *Pto* DC3000 (Gimenez-Ibanez et al. 2019). SA-JA antagonism during pathogen infections was also observed in *M. polymorpha* (Gimenez-Ibanez et al. 2019; Matsui et al. 2020). In *A. thaliana, coi1* mutants display an enhanced resistance to *Pto* DC3000, which depends on the SA pathway (Kloek et al. 2001). In contrast, resistance to *Pto* DC3000 was not affected in *M. polymorpha coi1* (Mp*coi1*) mutants (Gimenez-Ibanez et al. 2019).

Stable transformation of *M. polymorpha* can be achieved by *Agrobacterium*-mediated transformation methods (Ishizaki et al. 2008; Kubota et al. 2013; Tsuboyama and Kodama 2014; Tsuboyama et al. 2018; Tsuboyama-Tanaka and Kodama 2015; Tsuboyama-Tanaka et al. 2015), indicating that *Agrobacterium* can also be utilized for transient gene expression in *M. polymorpha*. Actually, Gimenez-Ibanez *et al*. firstly demonstrated that *Agrobacterium*-mediated transient gene expression is feasible in *M. polymorpha*, although expression profiles of introduced genes with this approach were obscure (Gimenez-Ibanez et al. 2019). Here, we demonstrate that reporter genes can be rapidly and robustly expressed in assimilatory filaments and rhizoids of *M. polymorpha* Tak-1 thalli simply by co-culturing with *A. tumefaciens* for a few days. We further show that our method is applicable in other *Marchantia* genotypes and species. Analysis of the Mp*coi1* mutant suggests that our method can be further improved by modulating phytohormone pathways, and can also be used as a novel pathosystem to address the roles of phytohormone pathways during *Marchantia*-bacteria interactions.

## Results

Sporelings, gemmae and regenerating thalli of *M. polymorpha* can be stably transformed using *Agrobacterium*-mediated methods (Ishizaki et al. 2008; Kubota et al. 2013; Tsuboyama and Kodama 2014; Tsuboyama-Tanaka and Kodama 2015; Tsuboyama-Tanaka et al. 2015). We therefore asked whether we can transiently express genes or proteins in *M. polymorpha* thalli simply by co-culturing with *A. tumefaciens* for short time periods. β-glucuronidase (GUS) reporters were used to monitor protein expression by GUS histochemical staining or by kinetic assays measuring GUS activity in thallus extracts. As *GUS* genes, we tested *uidA* from *Escherichia coli* and *GUSPlus*, a synthetic *gusA* gene from *Staphylococcus sp*. (Broothaerts et al. 2005). Additionally, we utilized the *GUSPlus* gene containing intron (*intron-GUSPlus*) to discriminate GUS activity that could potentially arise from *Agrobacterium* cells expressing *GUS* genes. The *GUS* genes were either driven by cauliflower mosaic virus (CaMV) 35S or *M. polymorpha EF1α* (Mp*EF1α*) promoters, which are frequently utilized in *M. polymorpha* for constitutive expression (Althoff et al. 2014).

Two-week-old *M. polymorpha* Tak-1 thalli, which were grown on agar plates, were transferred to liquid medium and co-cultured with *A. tumefaciens* strain GV3101::pMP90 harboring the GUS reporter plasmids, on a shaker (Fig. 1A - 1D). After 3 days of co-cultivation, GUS activity was visualized by histochemical staining. Notably, GUS staining was mainly observed in assimilatory filaments and rhizoids (Fig. 1E - 1H). Assimilatory filaments are located in air chambers of the photosynthetic layer at the thallus surface, and according to the distribution of air chambers a spotty pattern was observed. A similar staining pattern was observed with all tested constructs (Fig. 1M - 1O and 2A - 2D). These results suggest that *A. tumefaciens* preferentially targets and transfers T-DNA into cells of assimilatory filaments and rhizoids. Interestingly, GUS staining of rhizoids was enhanced when we used *uidA* compared to *GUSPlus* (Fig. 2C and 2D). We were able to rule out the possibility that the detected GUS activities were derived from *A. tumefaciens* cells that may have remained on the analyzed tissues, but not from infected *M. polymorpha* cells, because significant differences were not observed in GUS activities and GUS staining patterns when using either *GUSPlus* or *intron-GUSPlus* (Fig. 1E – 1O and Fig. S1). The use of two promoters, CaMV35S and Mp*EF1α*, gave similar results (Fig. 1M-1O and Fig. S1). Furthermore, we found that rhizoids of 0-day-old gemmae, which were harvested from gemma cups, can be used for the transient transformation with a certain probability (Fig. 1P).

**Figure 1.**
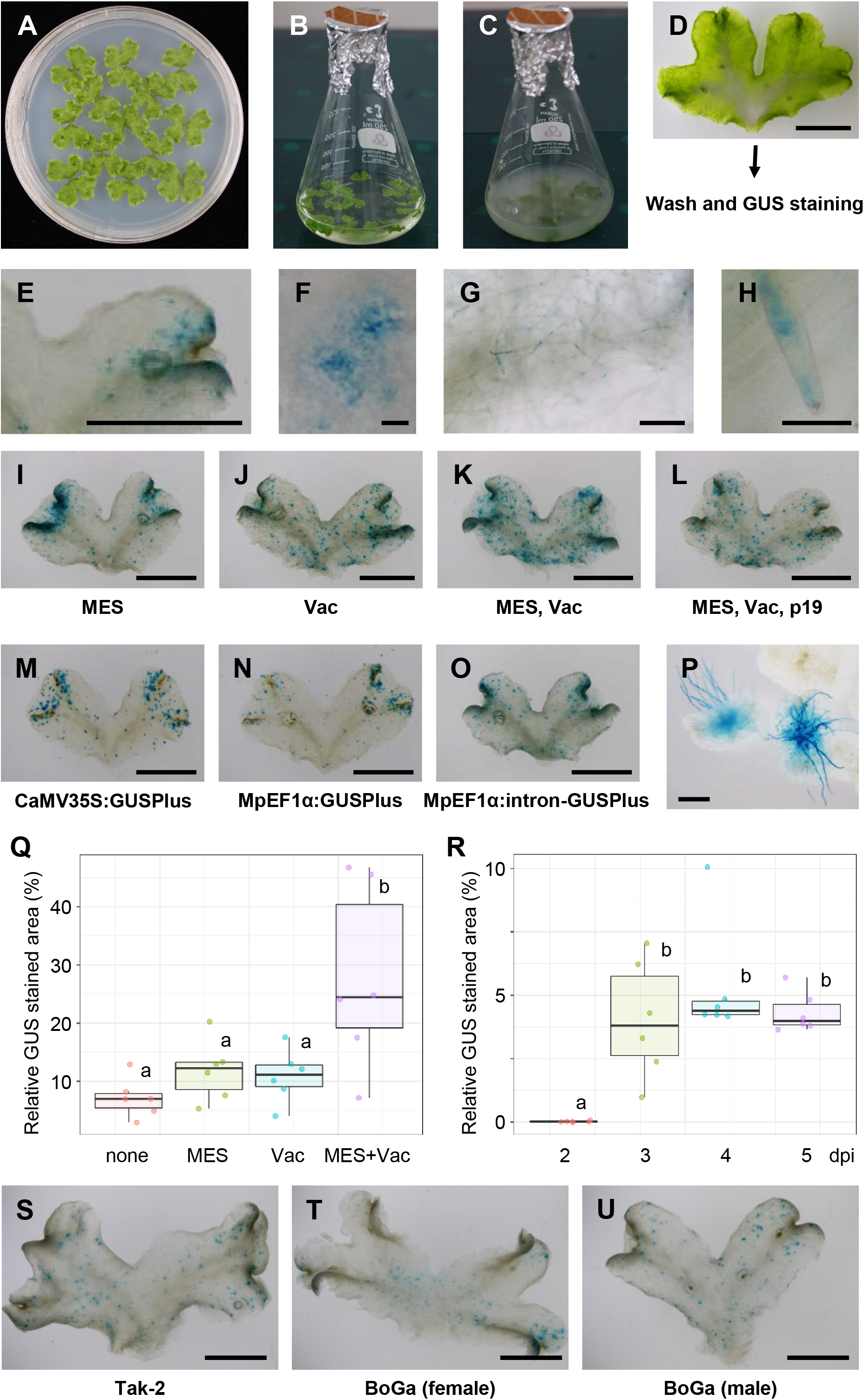
Transient *GUSPlus* gene expressions introduced by *Agrobacterium*-mediated transformation. (A - D) Procedures of *Agrobacterium*-mediated transient transformation into *M. polymorpha*. 14-day-old thallus of Tak-1 (A) and *Agrobacterium* harboring *proCaMV35S:intron-GUSPlus* construct are co-cultured. Initial and terminal (3 days) co-culture are shown in B and C, respectively. Agrobacterium are accumulating around rhizoids (D). (E - H) Histochemical GUS staining of Tak-1 thallus. Dorsal and ventral views are shown in E and G, respectively. Expanded images of air pore and rhizoids are shown in F and H, respectively. (I - L) Histochemical GUS staining of Tak-1 thallus that were transformed under different conditions as indicated, which are MES supplementation into co-culture medium (MES), vacuum infiltration before co-culture (Vac), and co-transformation with p19 (p19). (M - O) Histochemical GUS staining of Tak-1 thallus transformed with indicated constructs. (P) Histochemical GUS staining of Tak-1 gemmae. *Agrobacterium* harboring *proCaMV35S:intron-GUSPlus* and the co-culturing condition with MES and vacuum infiltration were used. (Q) Relative GUS stained areas of Tak-1 thalli transformed under different conditions. Different lower letters indicate a significant difference (Tukey’s test; P < 0.01). (R) Relative GUS stained areas of Tak-1 thalli transformed under different co-culture times. Different lower letters indicate a significant difference (Tukey’s test; P < 0.01). (S - U) Histochemical GUS staining of Tak-2 (S), female BoGa (T), and male BoGa (U). bars: 5 mm (D, E, I-O, S-U), 0.1 mm (F and H), and 0.5 mm (G and P)

**Figure 2.**
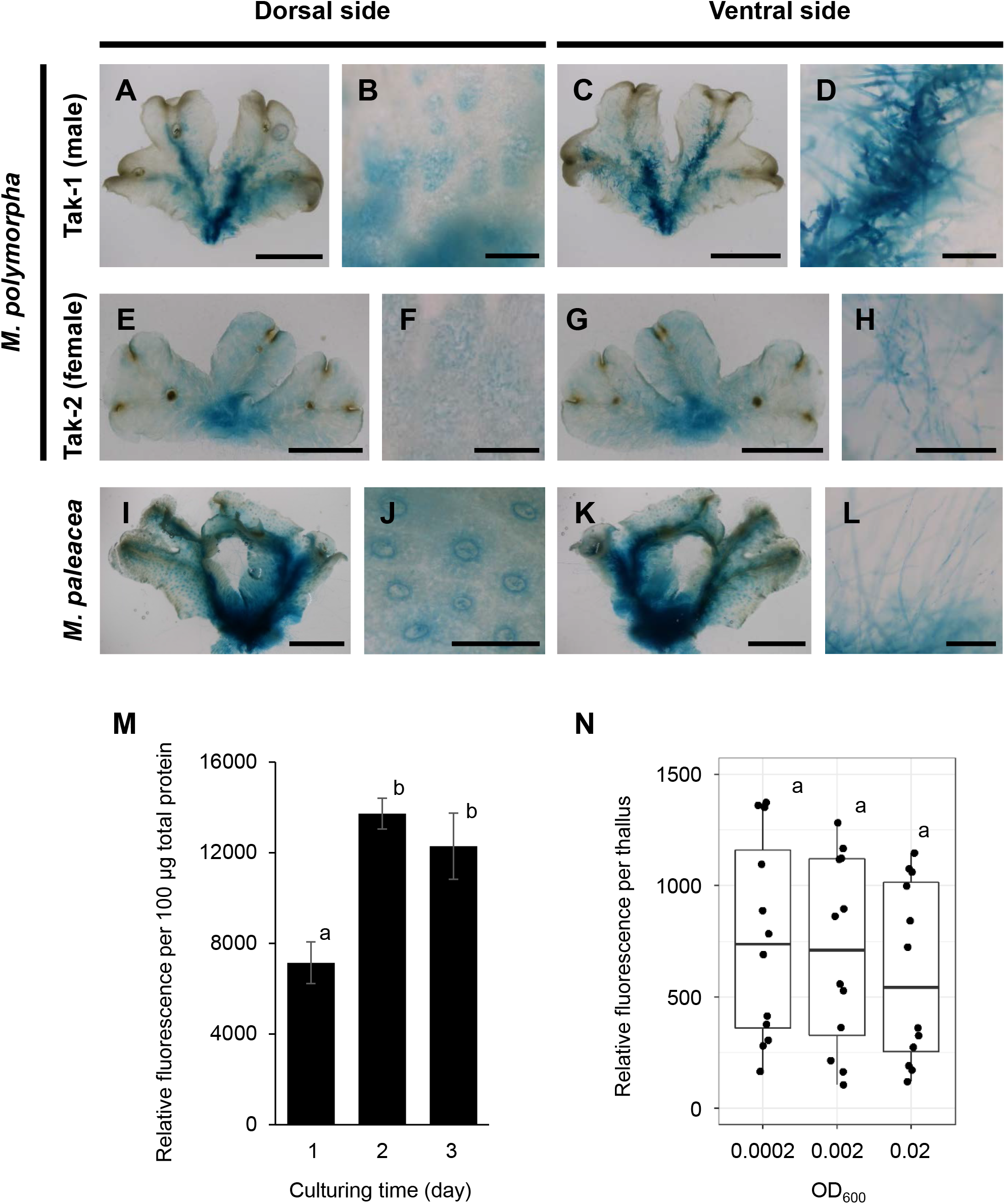
Transient *uidA* gene expressions introduced by *Agrobacterium*-mediated transformation. Histochemical GUS staining of Tak-1 (A - D), Tak-2 (E - H), and *M. paleacea* (I - L). Dorsal (A, E and I) and ventral (C, G, and K) views are shown. Expanded images of air pores (B, F, and J) and rhizoids (D, H, and L) are shown. bars: 5 mm (A, C, E, G, I, and K) and 0.5 mm (B, D, F, H, J, and L). (M) Relative GUS activity of Tak-1 thalli transformed under different co-culture times. Different lower letters indicate a significant difference (Tukey’s test; P < 0.01). (N) Relative GUS activity of Tak-1 thalli transformed under different initial bacterial concentration. Different lower letters indicate a significant difference (Tukey’s test; P < 0.01).

GUS staining revealed that assimilatory filaments in some, but not all, air chambers were successfully transformed. One plausible explanation for this observation is that *Agrobacterium* failed to enter all air chambers because of architecture and hydrophobicity of the air pore, which prevents the entry of water/liquid to air chambers to ensure efficient gas exchange (Schonherr and Ziegler 1975). Thus, applying vacuum is expected to assist *A. tumefaciens* to enter air chambers and thereby improve the transformation efficiency of assimilatory filaments. In *A. thaliana*, stabilizing pH at 5.5 with 2-(N-morpholino) ethanesulfonic (MES) buffer suppresses PTI and can enhance *Agrobacterium*-mediated transient gene expression (Wang et al. 2018). Therefore, we tested whether vacuum infiltration and/or pH stabilization with MES can boost the number of transformed cells by using *proCaMV35S:intron-GUSPlus*. As expected, vacuum infiltration resulted in more uniformly distributed GUS-stained spots throughout the thallus (Fig. 1I – 1K and Fig. S2). The pH stabilization did not affect the GUS staining pattern, but a trend towards an increased transformation efficiency was observed (Fig. 1E, 1I, and 1Q and Fig. S2). Strikingly, vacuum infiltration combined with pH stabilization could significantly improve the transformation efficiency (Fig. 1E, 1I - 1K, and 1Q and Fig. S2). We further tested whether co-expression of the RNA silencing suppressor p19 can enhance transient gene expression as in other plant species (Qiu et al. 2002; Qu and Morris 2002; Silhavy et al. 2002). Contrary to expectation, co-expression of p19 did not enhance but rather suppressed the GUS expression under the tested conditions (Fig. 1K and 1L and Fig. S2).

We further investigated the effect of co-cultivation duration on GUS expression. Expression levels were determined by measuring GUS-stained areas or GUS activity in transformed thallus extracts. When *proCaMV35S:intron-GUSPlus* was used, we could hardly detect GUS staining after 2 days of co-cultivation. The GUS staining became apparent and saturated after 3 days of co-cultivation (Fig. 1R and Fig. S3). Meanwhile, when we used *pro*Mp*EF1α:uidA*, GUS activity was detected after 1 day of co-cultivation and saturated after 2 days of co-cultivation (Fig. 2M). The observed differences could be due to an increased expression of *uidA* in rhizoids or usage of different measures for detecting GUS activity. Based on the measured GUS activity by using *pro*Mp*EF1α:uidA*, the initial dose of *A. tumefaciens* for co-cultivation in the range of OD_600_ 0.0002-0.02 is likely to have no influence on the transformation efficiency (Fig. 2N).

To assess whether our *Agrobacterium*-mediated transient expression approach can be utilized for subcellular localization analysis, we expressed a fluorescently tagged protein, tdTomato, fused to a nuclear localization signal (tdTomato-NLS) (Ishizaki et al. 2015). As shown in Fig. 3A, we observed a fluorescent signal for tdTomato-NLS in the nucleus of an assimilatory filament cell as expected. We then attempted to co-express two fluorescently tagged proteins; Citrine-MpNPSN1, which localizes at the plasma membrane (Kanazawa et al. 2016), and tdTomato-NLS. As shown in Fig. 3B, we observed co-expression of the two fluorescently tagged proteins in the same cell, in which the expressed proteins displayed the expected localizations. This suggests that co-localization or co-immunoprecipitation assays could possibly be performed to investigate protein-protein interactions.

**Figure 3.**
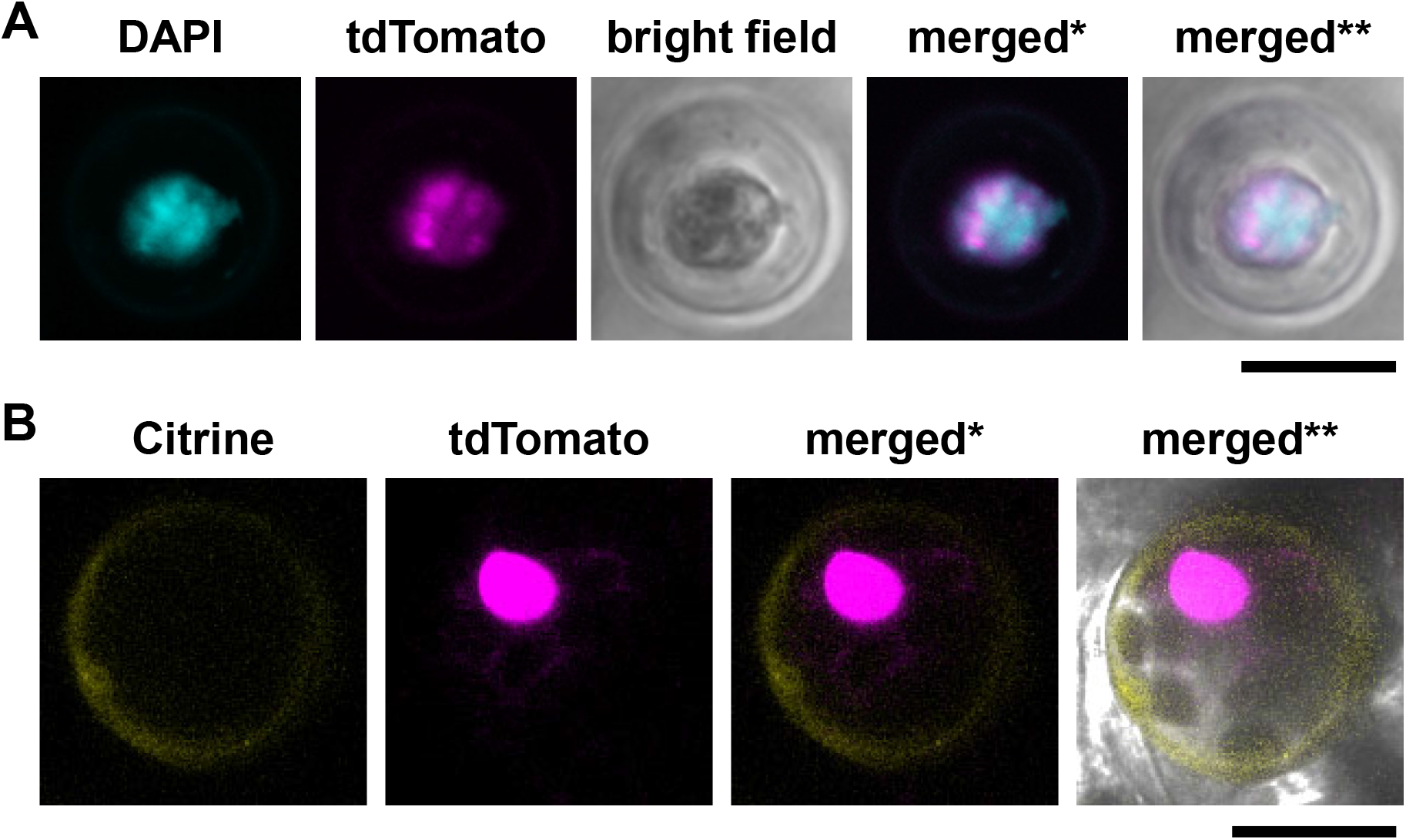
Subcellular localization of fluorescently tagged proteins. (A) Subcellular localization of tdTomato-NLS. Fluorescence from DAPI, tdTomato, and bright field images are shown. *: merged image of DAPI and tdTomato. **: merged image of DAPI, tdTomato, and bright field. (B) Subcellular localization of Citrine-MpNPSN1 and tdTomato-NLS. Fluorescence from Citrine, tdTomato, and bright field images are shown. *: merged image of Citrine and tdTomato. **: merged image of Citrine, tdTomato, and bright field. bars: 10 µm.

*M. polymorpha* accessions other than *M. polymorpha* Tak-1 have also been widely used for *Marchantia* research. Therefore, we assessed whether our method is applicable to other *M. polymorpha* accessions, Tak-2 and BoGa (Buschmann et al. 2016). As shown in Fig. 1S - 1U, GUS staining was observed in these accessions and the staining patterns were similar to what we observed in *M. polymorpha* Tak-1. Moreover, our method was also successful for other *Marchantia* species having symbiotic ability, *M. paleacea* (Fig. 2I - 2L) (Buschmann et al. 2016; Field et al. 2012). These results suggest that our method can be broadly applied to *M. polymorpha* and to other non-model liverworts.

We next explored whether the above demonstrated *Agrobacterium*-mediated GUS expression method can be utilized as a novel tool or pathosystem to study liverwort-bacteria interactions. In *M. polymorpha*, the JA receptor mutants, Mp*coi1-1* and Mp*coi1-2*, were shown to display the same levels of resistance against the bacterial pathogen *Pto* DC3000 compared to wild-type Tak-1 and Tak-2 (Gimenez-Ibanez et al. 2019). Thus, it is not suitable to utilize *Pto* DC3000 for investigating contributions of JA in *M. polymorpha*. In *A. thaliana*, it is known that *Agrobacterium*-mediated transient transformation is restricted in the At*coi1* mutant, lacking a JA receptor (Rosas-Díaz et al. 2017). Therefore, we investigated whether the *Agrobacterium*-mediated *intron-GUSPlus* expression is altered in the Mp*coi1-2* mutant (Monte et al. 2018). As expected, detected GUS activity in the Mp*coi1-2* mutant was significantly lower compared to that in wild-type Tak-1 (Fig. 4A – 4C, Fig. S5). This result suggests that JA signaling pathway contributes to resistance against *A. tumefaciens* in *M. polymorpha* as in *A. thaliana*, and secures the relevance of this method to study JA pathway in liverwort-bacteria interactions.

**Figure 4.**
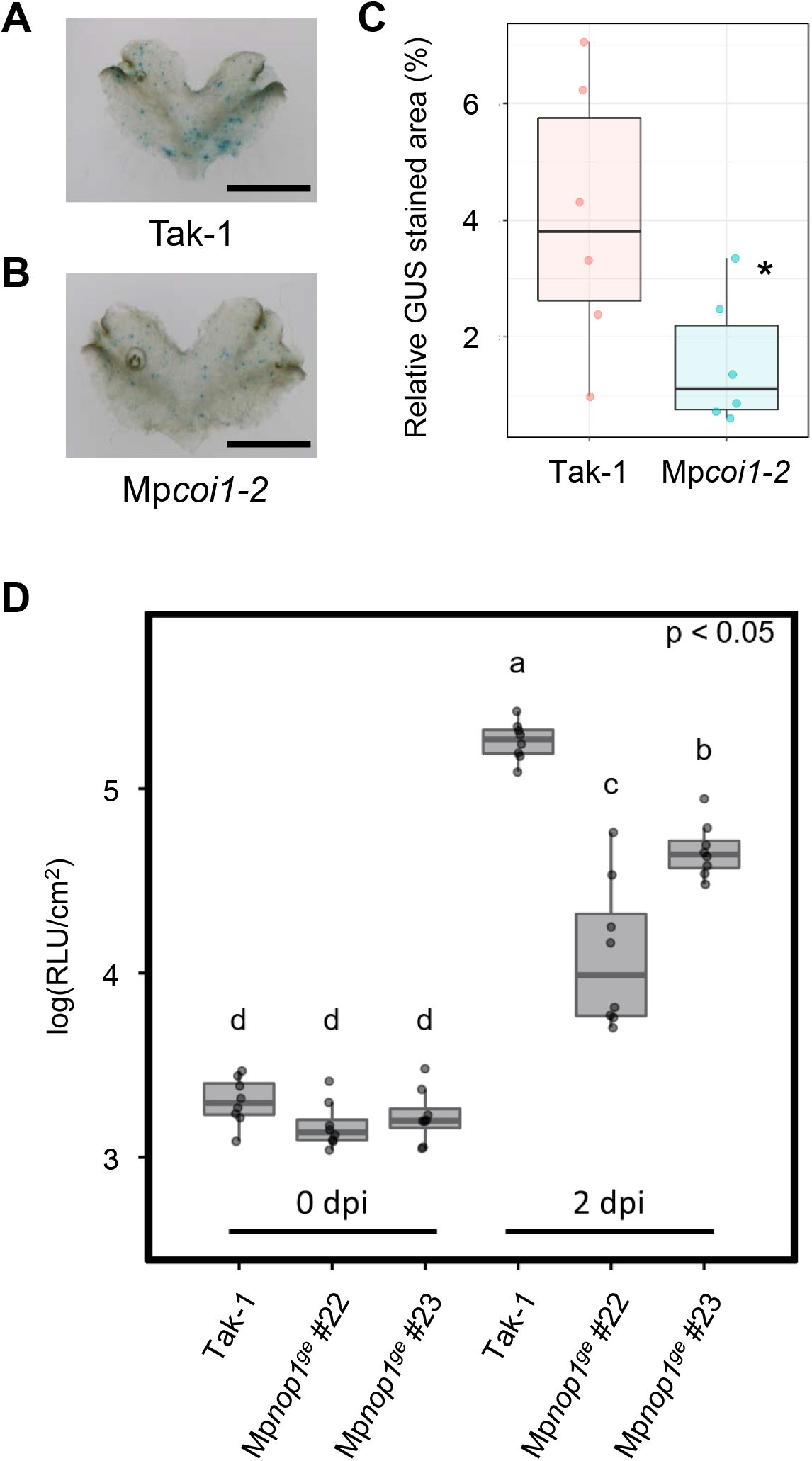
(A and B) Histochemical GUS staining of Tak-1 (A) and Mp*coi1-2* (B) transformed with *proCaMV 35S:intron-GUSPlus*. (C) Relative GUS stained areas of Tak -1 and Mp*coi1-2* thalli. bars: 5 mm. (D) Fourteen-days-old thalli of Tak-1 and Mp*nop*^*ge*^ mutant lines were vacuum infiltrated with *Pto*-lux at OD_600_ = 0.01. Log10-transformed RLU were calculated from 8 biological replicates collected from thalli of different plants. Statistically significant differences are indicated by different letters (adjusted P < 0.05).

The bacterial pathogen *Pto* DC3000 is able to infect and cause disease on *M. polymorpha* in an effector-dependent manner (Gimenez-Ibanez et al. 2019; Matsumoto et al. 2021). However, its infection strategy has yet to be clarified. Gimenez-Ibanez *et al*. demonstrated that *Agrobacterium*-mediated transient expression of *AvrPto* or *AvrPtoB* in *M. polymorpha* thalli could suppress Mp*CML42*, Mp*WRKY22*, and Mp*ACRE132* gene expression induced by crude extracts from *Pto* DC3000 (Gimenez-Ibanez et al. 2019). Together with our observation that *A. tumefaciens* preferentially targets assimilatory filaments that are located in air chambers, we hypothesized that *Pto* DC3000 colonization is supported by entry into air chambers and targeting assimilatory filaments, as in the case of infection of *M. polymorpha* with the oomycete pathogen *Phytophthora palmivora* (Carella et al. 2018). To address this possibility, we inoculated the air-chamberless *nop1* mutants, Mp*nop1*^*ge*^, with the bioluminescent *Pto* DC3000 (*Pto*-lux) (Ishizaki et al. 2013; Matsumoto et al. 2021; Shimamura 2016a; Sugano et al. 2018). As expected, *Pto*-lux growth in the Mp*nop1* mutants was significantly reduced compared to the growth in wild-type Tak-1 (Fig. 4D). This result implies that the air chamber is an initial battlefield for bacterial pathogens to successfully colonize liverworts.

## Discussion

In this report, we described a simple and robust *Agrobacterium*-mediated transient transformation method and its performance in expressing genes in *Marchantia* thalli. We observed that *Agrobacterium* primarily and efficiently transferred the *GUS* reporter genes into rhizoids and assimilatory filaments, which are specialized cell types for water/nutrient transport and photosynthesis, respectively (Cao et al. 2014). This observation is reasonable as it is known that these cell types are less covered by cuticles that play important roles in plant defense against diverse pathogens (Schonherr and Ziegler 1975). Our results may suggest that assimilatory filaments and rhizoids are the primary site at which microbes interact with liverworts. In support of this hypothesis, reports have demonstrated that the hemi-biotrophic oomycete pathogen *Phytophthora palmivora* establishes successful infection by colonizing air chambers (Carella et al. 2018), and that symbiotic fungi enter through rhizoids (Humphreys et al. 2010). Along the same lines, we revealed that air chambers support colonization of the bacterial pathogen *Pto* DC3000 (Fig. 4D).

Transient gene expression in epidermal cells remained a major challenge. The enhanced transformation efficiency by pH stabilization implies that *M. polymorpha* can recognize *A. tumefaciens* through MAMP receptors, as is the case for *A. thaliana*. Thus, by gaining a deeper understanding of the immune system in *M. polymorpha* or in other liverworts, further method improvement can be achieved. Since we revealed that the Mp*coi1-2* mutant is resistant to *A. tumefaciens* infection, it would be interesting to test SA pathway-deficient *M. polymorpha* mutants, although we need to begin with understanding the biological functions of the SA and SA pathway in *M. polymorpha*.

Expression analysis of the fluorescently tagged proteins confirmed that our method can be used to study the subcellular localization of proteins. In addition, the use of two kinds of *Agrobacterium* cells with different expression vectors allowed observation of two proteins from each vector in the same cell (Fig. 3B), which has also been reported for stable transformation events (Ishizaki et al. 2015). Thus, our transient expression method can be exploited to quickly analyze co-localization of different proteins and protein-protein interactions. However, it should be noted that expressed proteins may mis-localize because of overexpression or unnatural conditions caused by the *Agrobacterium* infection procedure, as is the case of transient expression in other plant species. Indeed, we did observe unusual localization of MpNEK1-Citrine in rhizoids, as shown in Fig. S4 (Otani et al. 2018).

Co-culturing is a key element of our method, because T-DNA is transferred and gene expression is initiated during this step. Our observation that GUS expression was detected within 2 days of co-cultivation, and saturated at 2 to 3 days, is in line with the effect of co-culture periods on stable transformation efficiency in *M. polymorpha* (Ishizaki et al. 2008; Tsuboyama and Kodama 2014). The initial concentration of co-cultivated *Agrobacterium* had no significant effect on transient GUS expression within the tested OD_600_ range of 0.0002-0.02, which is slightly different from stable transformation with the AgarTrap method whose efficiency drops below an OD_600_ of 0.3 (Tsuboyama-Tanaka and Kodama 2015). Our *Agrobacterium*-mediated transient transformation method is likely to be more robust in this sense.

The finding that our method is applicable to *M. paleacea* opens up exciting new possibilities. For instance, as *M. paleacea* and other liverworts initiate symbiotic interactions with fungi at rhizoids, our method for efficient transformation of rhizoids could be a useful tool for understanding the molecular mechanisms of these plant-fungal interactions. It might also be possible to combine our method with a transient gene silencing technique, such as virus-induced gene silencing (VIGS). Moreover, as the body architecture of different liverwort species exhibit many similar features, our method could be applied to liverworts other than *Marchantia*.

## Materials and Methods

### Plant material and growth conditions

*M. polymorpha* accessions Tak-1, Tak-2, BoGa (Althoff et al. 2014; Bowman et al. 2017; Buschmann et al. 2016), and *M. paleacea* (Field et al. 2012) were used in this study. Plants were grown on half-strength Gamborg’s B5 basal salt mixture medium (Duchefa, Netherlands) containing 1 % plant agar (Duchefa, Netherlands) at 22 °C, under continuous light or 14 h light/10 h dark cycles (50 µmol photons m^-2^s^-1^ white LED).

### Plasmid construction

The binary plasmids pMpGWB302-GUSPlus, pMpGWB302-intron-GUSPlus, pMpGWB303-GUSPlus, pMpGWB303-intron-GUSPlus, and pMpGWB303-GUS harboring *proCaMV35S:GUSPlus, proCaMV35S:intron-GUSPlus, pro*Mp*EF1α:GUSPlus, pro*Mp*EF1α:intron-GUSPlus*, and *pro*Mp*EF1α:uidA*, respectively, were constructed by using the gateway system. The pMP22, pUL22, and pENTR-gus (ThermoFisher U.S.A.) harboring *GUSPlus, intron-GUSPlus*, and *uidA*, respectively, were mixed with pMpGWB302 and pMpGWB303 harboring CaMV35S promoter (*proCaMV35S*) and a promoter of Mp*EF1α* (*pro*Mp*EF1α*), respectively (Ishizaki et al. 2015), and the LR reaction was performed. The binary plasmid pMpGWB116-proMpEFα harboring *pro*Mp*EF1α:tdTomato-NLS* was constructed in the same way using pENTR-proMpEF1α and pMpGWB116 (Ishizaki et al. 2015). The binary plasmid pMpGWB305-Citrine-MpNPSN1 harboring *proCaMV35S: Citrine-*Mp*NPSN1* was described previously (Kanazawa et al. 2016). The plasmid pBICp19 harboring *proCaMV35S:p19* was described previously (Takeda et al. 2002).

### Agrobacterium-mediated transient transformation

To prepare *Agrobacterium* for infection, one of the binary plasmids or pBICp19 was transformed into *Agrobacterium tumefaciens* strain GV3101::pMP90. A single colony of bacteria was inoculated into Luria-Bertani (LB) medium containing spectinomycin antibiotics and cultured for two days at 28 °C. Bacterial cells were collected from 1 ml culture by centrifugation, suspended in 5 ml 0M51C medium (Ono et al. 1979; Takenaka et al. 2000) containing 2 % sucrose and 100 µM 3,5-dimethoxy-4-hydroxyacetophenone (acetosyringone), and cultured for 5 h at 28 °C. In parallel, *M. polymorpha* (Tak-1, Tak-2, and BoGa) and *M. paleacea* gemmae were cultured on the agar plates for 14 or 18 days at 22 °C under continuous light or a 14 h light/10 h dark cycle. The bacterial culture was diluted with 0M51C containing 2 % sucrose and 100 µM acetosyringone to be at OD_600_ = 0.02 or values specified in Fig. 2F and then 10 *Marchantia* thalli were transferred into the 50 ml bacterial suspension. The *Marchantia* thalli and *Agrobacterium* were co-cultured at 22 °C under a 14 h light/10 h dark cycle for 1 to 5 days with shaking (120 rpm). If not specified, plants were co-cultured with *Agrobacterium* for 3 days. To test the effect of p19, bacteria transformed with the binary plasmid or pBICp19 were mixed in a 1:1 ratio and used for the co-culturing.

### Histochemical GUS staining

Histochemical assays for GUS activity were performed according to a previously described procedure (Jefferson et al. 1987), with some modifications. Briefly, after the co-culturing, *Marchantia* thalli were washed 5 times with water, fixed in 90 % acetone at -20 °C for 1 to 5 days, washed twice with water, and incubated in GUS staining solution containing 1 mM 5-bromo-4-chloro-3-indolyl-β-D-glucuronic acid (X-Gluc: X-Gluc DIRECT, U.K.), 50 mM sodium phosphate pH 7.0, 1 mM EDTA, 0.1 % Triton X-100, 10 mM β-mercaptoethanol at 37 °C for 8-16 hours. To enhance the contrast of GUS staining, chlorophyll was removed with 70 % (v/v) ethanol.

### Quantification of GUS activity

The quantitative GUS assay was performed as described previously (Halder and Kombrink 2015), with some modifications. *Marchantia* thalli were washed five times with water after the co-culturing, transferred to microtubes, milled in the extraction buffer containing 100 mM sodium phosphate pH 7.0, 1 mM DTT, and centrifuged at 25,000 rcf, 4 °C for 30 min. The clear supernatant (50 µl) was transferred to a 96-well transparent plate, mixed with GUS assay solution (50 µl) containing 2 mM 4-methylumbelliferyl-β-D-glucuronide (4-MUG), 50 mM sodium phosphate pH 7.0, 1 mM EDTA, 0.1 % Triton X-100, and 10 mM β-mercaptoethanol. Samples were incubated at 37 °C for 120 min and 4-MU fluorescence was measured every 10 min using the FLUOstar Omega microplate reader (BMG LABTECH, Germany) with an excitation/emission wavelength of 355/460 nm. GUS activity was calculated using the ΔE_460_ increments (10-120 min).

### Confocal laser scanning microscopy

For fluorescence microscopy analysis with 4’,6-diamidino-2-phenylindole (DAPI) staining, samples were fixed with fixation buffer containing 4 % paraformaldehyde, 0.1 M sodium phosphate, pH 7.0, washed twice with water, incubated in 1 µg/ml DAPI solution, and mounted in water and observed using an LSM780 (Carl Zeiss, Germany) equipped with a water immersion lens. The samples were excited at 405 nm for DAPI, and 561 nm for tdTomato, and emission was collected between 410 nm and 585 nm, and between 580 nm and 650 nm, respectively. The images were processed digitally with Zen2011 (Carl Zeiss, Germany) and Photoshop (Adobe systems, U. S. A). To analyze the fluorescence of cells that co-transformed MpNEK1-Citrin and TagRFP-MpTUB2, samples were mounted without fixation in water and observed using an LSM780. The samples were excited at 514 nm for mCitrine, and 561 nm for TagRFP, and emission was collected between 519 nm and 574 nm, and between 580 nm and 650 nm, respectively.

### Pto DC3000 growth assay

The growth of *Pto* DC3000 was measured by using the *Pto* P_kan_:lux strain as previously described (Matsumoto et al. 2021). *M. polymorpha* Tak-1 gemmae were placed on the half-strength Gamborg’s B5 agar covered with a Whatman filter paper (Cat. No. 1001-085) and grown for 14 days at 22 °C under continuous white LED light. The 14-day-old thalli were transferred to fresh empty Petri dishes, submerged in the *Pto* DC3000 suspensions in sterile water at OD_600_ = 0.01, and incubated with vacuum for 5 min. After the incubations, thalli were placed onto a Whatman filter paper, which had been soaked with Milli-Q water in fresh empty Petri dishes. Inoculated thalli were kept in a climate chamber at 22 °C under a 16 h light/8 h dark cycle and sampled at 0 and 2 dpi. One biological replicate consisted of single thallus disc (5 mm diameter) that was excised from the basal region of individual thallus using a biopsy punch, and 8 biological replicates were collected. The thallus discs were transferred to wells of a white reflecting 96-well plate (VWR, #738-0016). The plate was placed in Fluostar Omega (BMG Labtech) plate reader and kept in the dark for 10 min before measurement to reduce background signals. Then, luminescence was measured for 5 s for each sample. Log10-transformed relative luminescence units (RLU) was calculated. Pairwise comparison was performed using the R function pairwise.t.test with pooled SD, and the Benjamini-Hochberg method was used for correcting the multiple hypothesis testing. Linear regression and calculation of correlation coefficients were performed using the R function lm and cor, respectively.

## Acknowledgements

The authors are grateful to Takayuki Kohchi (Kyoto University, Japan) for providing pMpGWB302, pMpGWB303, and Mp*nop1*^*ge*^ mutants, Ivan F. Acosta (MPIPZ, Germany) for providing pMP22 and pUL22, Takeshi Ueda (NIBB, Japan) for providing pMpGWB305-Citrine-MpNPSN1, Atsushi Takeda (Ritsumeikan University, Japan) for providing pBICp19, Kenichi Tsuda (MPIPZ, Germany) for providing *A. tumefaciens* GV3101::pMP90, Sabine Zachgo (Osnabrück University, Germany) for providing *M. polymorpha* accession BoGa, and Sebastian Schornack (University of Cambridge, U.K.) for providing *M. paleacea*. We also thank Rozina Kardakaris and Neysan Donnelly (MPIPZ, Germany) for editing the manuscript.

## Legends for supplementary figures

**Figure S1** Relative GUS activity of Tak-1 thalli transformed with different constructs. Different lower letters indicate a significant difference (Tukey’s test; P < 0.01).

**Figure S2** (A - E) Images of *Marchantia* thalli used for statistical analysis shown in Fig. 1Q. Conditions of transformation are shown in each panel. bars: 5 mm. (F) Relative GUS stained areas of Tak-1 thalli transformed under different conditions. Different lower letters indicate a significant difference (Tukey’s test; P < 0.01).

**Figure S3** (A - D) Images of *Marchantia* thalli used for statistical analysis shown in Fig. 1R. Co-culturing times (days) are shown in each panel. bars: 5 mm.

**Figure S4** (a) Co-expression of fluorescently tagged proteins. *pro*Mp*NEK1:* Mp*NEK1-Citrine* and *proCaMV35S:TagRFP-*Mp*TUB2* were co-transformed into Tak-1. Fluorescence from Citrine and TagRFP and merged images are shown. bar: 50 µm.

**Figure S5** (A and B) Images of *Marchantia* thalli used for statistical analysis shown in Fig. 4C. Genotypes are shown in each panel. bars: 5 mm.

